# A biologically driven directional change in susceptibility to global-scale glaciation during the Precambrian-Cambrian transition

**DOI:** 10.1101/359422

**Authors:** Richard A. Boyle, Carolin R. Löscher

## Abstract

Integrated geological evidence suggests that grounded ice sheets occurred at sea level across all latitudes during two intervals within the Neoproterozoic era; the “snowball Earth” (SBE) events. Glacial events at ~730 and ~650 million years ago (Ma) were probably followed by a less severe but nonetheless global-scale glaciation at ~580Ma, immediately preceding the proliferation of the first fossils exhibiting unambiguous animal-like form. Existing modelling identifies weathering-induced *CO*_2_ draw-down as a critical aspect of glacial inception, but ultimately attributes the SBE phenomenon to unusual tectonic boundary conditions. Here we suggest that the evident directional decrease in Earth’s susceptibility to a SBE suggests that such a-directional abiotic factors are an insufficient explanation for the lack of SBE events since ~580 Ma. Instead we hypothesize that the terrestrial biosphere’s capacity to sustain a given level of biotic weathering-enhancement under suboptimal/declining temperatures, itself decreased over time: because lichens (with a relatively robust tolerance of sub-optimal temperatures) were gradually displaced on the land surface by more complex photosynthetic life (with a narrower temperature window for growth). We use a simple modelling exercise to highlight the critical (but neglected) importance of the temperature sensitivity of the biotic weathering enhancement factor and discuss the likely values of key parameters in relation to both experiments and the results of complex climate models. We show how the terrestrial biosphere’s capacity to sustain a given level of silicate-weathering-induced *CO*_2_ draw-down is critical to the temperature/greenhouse forcing at which SBE initiation is conceivable. We do not dispute the importance of low degassing rate and other tectonic factors, but propose that the unique feature of the Neoproterozoic was biology’s capacity to tip the system over the edge into a runaway ice-albedo feedback; compensating for the self-limiting decline in weathering rate during the temperature decrease on the approach to glaciation. Such compensation was more significant in the Neoproterozoic than the Phanerozoic due, ultimately, to changes in the species composition of the weathering interface over the course of evolutionary time.

## 1. Introduction

### A. The snowball Earth problem

The snowball Earth (SBE) concept provides a rich framework for quantifying the limits of integrated biogeochemical change (Hoffman & Schrag, 2002). The period between 720-635 Ma is termed the Cryogenian (from Greek words for “cold” and “birth”) and an iconic feature of this interval is the presence of radiometrically dated glacial diamictites in over ninety formations worldwide (Evans, 2000, Arnaud et al, 2012, Hoffman et al, 2017). Many such deposits exhibit both the appearance of derivation from glaciers grounded below sea-level and a remnant magnetism indicative of a low paleolatitude (Evans & Raub, 2011). This is widely taken to imply that globally extensive glaciers progressed as far as the equator. Numerous other diamictites are bounded by thick layers of non-skeletal shallow-water carbonate (Fairchild, 1993, Hoffman & Halverson, 2008) likely derived from the warmest parts of the ocean (Opdyke & Wilkinson, 1993). Equivalently, this is taken to imply that the glacial interval’s termination was characterized by an extreme *CO*_2_ greenhouse, which gradually declined as the ocean “titrated” the *CO*_2_ as carbonate. This core idea motivated the original Snowball Earth hypothesis (Kirshvink, 1992, Hoffman et al, 1998) and has remained relatively robust to subsequent scrutiny. Contemporary discussion as to how best to interpret the data motivating SBE centres around the relevance of incompletely oxidized iron-rich rocks (Urban et al, 1992, MacDonald et al, 2010), other evidence pointing to the redox status of the ocean during the glacial interval (Halverson, 2011) and the plausibility of the persistence of habitable refugia for eukaryotes (Hoffman, 2016).

A key conceptual uncertainty within the SBE concept is the degree of dynamism present during the glacial interval. A “slushball-Earth” is a global-scale glaciation with open ocean water towards equatorial latitudes, because an extra (greenhouse) forcing causes the ice-albedo feedback to stabilize dynamically, rather than runaway to an equilibrium, entailing complete ice cover (e.g. Peltier, 2007). The concept of a slushball has historically been defined in opposition, and widely held to be more parsimonious than, a “hard snowball” in which the ice-albedo feedback goes to an equilibrium state. This is because most researchers have tacitly assumed that a slushball is far easier to reconcile with the survival of complex life, and/or that a hard snowball would lead to accumulation of ice sheets too thick to permit deglaciation. Thin ice solutions within global climate models are indeed extremely precarious (Pollard & Kasting, 2005, Lewis et al, 2007), generally hinging on the action of some process within the hydrological cycle (evapotranspiration, dynamic sea ice, wind fields) that can transport ice from the tropics poleward, thereby avoiding the accumulation of irretrievably thick ice sheets at the equator. However, high-pressure-induced melting at the base of extremely thick ice sheets may induce “basal sliding”, potentially sustaining a sufficiently dynamic sea ice cycle to permit deglaciation from a hard snowball (Hoffman, 2005, Rignew et al, 2011). Hard snowball solutions within general circulation models can be formulated that permit deglaciation, but this is by no means a foregone conclusion (Abott et al 2013). Again, the reversibility of the hard-snowball state almost always hinges on crucial unpreserved quantities, such as tropical evaporation fluxes (Pollard & Kasting, 2005), heat transfer patterns within the atmosphere above an ice-covered ocean (Benn et al, 2015), or the glacial interval’s cloud formation patterns and their contribution to albedo (Abbot et al, 2012). This raises the (pessimistic but very real) possibility that the data do not allow us to distinguish between a hard snowball and slushball solution. Nevertheless, the broad view that emerges from current work with complex climate models is that a hard snowball, with sufficient oceanographic/atmospheric dynamism to keep equatorial ice recoverably thin, is probably closest to the mark (reviewed in e.g. Hoffman, 2018).

In this context the utility of the SBE problem as a theoretical test case for the predictive power of complex climate models becomes apparent, as does the fact that the “hard-snowball versus slushball” distinction cannot be resolved in a hypothesis-driven simple-modelling study such as this. We therefore proceed by remaining agnostic about this distinction. We simply take the label “snowball Earth” to refer to the entry of Earth into a positive feedback between low-latitude ice cover and high planetary albedo, with the result that some form of equilibrium is reached. (In a hard-snowball this would constitute a truer equilibrium in an “energy balance” sense, in a slushball the “equilibrium” might be better described as a dynamical steady state. The key point is that the ice-albedo feedback reaches a region of “phase space” qualitatively distinct from subsequent milder glaciations).

### B. The silicate weathering CO_2_-sink, climate stability and the ice-albedo feedback

Weathering of silicates on the land surface removes *CO*_2_ from the atmosphere and inputs bicarbonate anions and silicate-derived (magnesium and calcium) cations into the water table, ultimately leading to precipitation of calcite on the ocean floor in a net *CO*_2_-sink (e.g. Berner et al, 1983):

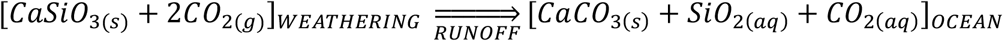

Because the weathering reaction is both temperature-sensitive and a greenhouse gas sink, a silicate weathering “thermostat” was postulated (Walker et al, 1981) to exert a homeostatic influence on Earth’s temperature over geologic timescales. A shutdown of this weathering sink during the glacial interval of a snowball Earth is central to the coherence of the original idea (Kirschvink, 1992), because without the resultant *CO*_2_ build-up, the combination of high albedo and low temperature would render global-scale ice cover irreversible (e.g. Pierrehumbert, 2005). Existence of a climatically influential weathering-homeostat presupposes kinetic limitation of weathering rate, at least at first-order, by *CO*_2_ levels. The idea of such limitation is therefore complicated by the fact that weathering rate may in fact frequently be limited by supply of fresh rock from tectonic uplift (West et al, 2005). Consistently, supply rate of silicate rock has been proposed to have controlled the interval between snowball Earth events- because deglaciation requires a high *CO*_2_ greenhouse, which must be removed by the weathering sink before the system becomes susceptible to a subsequent glacial event (Mills et al, 2011).

An ice-covered surface reflects a higher fraction of incoming radiation (has a higher albedo) than an ice-free surface, which leads to a hysteresis (memory of previous system state) in the climate system, by which the greenhouse/radiative forcing necessary to exit the snowball state exceeds the threshold at which the system initially fell into this state (Budyko, 1969, Sellers, 1969). If more than about half of Earth’s surface becomes ice-covered, a runaway positive feedback to global glaciation becomes inevitable (Caldeira & Kasting, 1992b). But the precise temperature/greenhouse forcing at which this albedo threshold occurs is non-trivial to calculate even in high-resolution climate models, because it depends on paleogeographic reconstructions, the contribution of cloud albedo, the albedo of ice-free but extremely low temperature ocean water the thickness of sea ice (Warren et al, 2002) and the density of cracks/bubbles in such sea ice (Carns et al, 2016), etc. Thus, it is possible to state that there exists a temperature *T*_*in*_ at which a positive feedback between declining temperatures, increasing ice-cover and increasing planetary albedo, will initiate, but the specific value of this temperature threshold is uncertain. Existing studies have taken *T*_*in*_ ≈ 10°C (Donnadieu et al, 2004, Pierrehumbert et al, 2005, Mills et al, 2011). Below, we parameterize a wider range of values for this threshold, on the basis of uncertainties in the appropriate inputs to the complex climate models from which this value was calculated, as well as uncertainty about what the snowball state actually constitutes in terms of temperature, albedo and greenhouse forcing combination.

### C. Biological enhancement of weathering, the evolution of the terrestrial biosphere and the plausibility of Neoproterozoic lichens

The biotic enhancement of weathering (BEW) factor is defined as how much faster the silicate weathering *CO*_2_ sink is under biotic conditions than under abiotic conditions (Schwartzman & Volk, 1989, Schwartzman, 2017). The magnitude of the BEW factor depends on the rock substrate and the principal form of photosynthetic life responsible for the enhancement. Empirical estimates suggest that for vascular plants *BEW* ≈ 2 − 5 on basalt (Moulton & Berner, 1998) and *BEW* ≈ 10 − 18 on granite (Bormann et al, 1998), for mosses *BEW* ≈ 1.4 − 3.5 on granite and *BEW* ≈ 3.6 − 5.5 on andesite (Lenton et al, 2012), for lichens *BEW* ≈ 4 − 16 on mica schist (Aghamiri & Schwartzman, 2002), *BEW* ≈ 2.5 − 3.5 on granite (Zambell et al, 2012) and probably of the order of *BEW* ≥ 15 − 20 (with some estimates suggesting significantly higher, up to 70) on basalt (Jackson & Keller, 1970, Brady et al, 1999, Stretch & Viles, 2002). Given the extreme uncertainties in the species composition by biomass of any ancient terrestrial biosphere, as well as in the rock composition of ancient paleo-continents, it is not realistic to formulate a detailed picture of the evolution of the *BEW* factor over time (although see Schwartzman, 2017). For the purposes of this exercise, we simply note that any enhancement of silicate weathering by lichens is comparable to, and if anything, greater than, that of more morphologically complex terrestrial autotrophs.

Mechanistically, lichens enhance weathering by proton secretion, ice-crystal nucleation and retention of water in the rock surface, secretion of organic acids, and digestion of silicate particles within the fungal hyphae (Jones & Wilson, 1985, Chen et al, 2000, Li et al, 2016). The adaptive value of this activity is likely the acquisition of the *PO*_4_^3−^ liberated with the silicate cations (Rogers, 1998, Guidry & McKenzie, 2000). Indeed, the rise of a Neoproterozoic biosphere dominated by lichens has been hypothesized previously, as the explanation for a possible rise in *O*_2_ at the Neoproterozoic-Cambrian boundary (Heckman et al, 2001, Lenton & Watson, 2004).

The first unambiguous fossil lichen is not observed until ~400Ma (Taylor et al, 1995), but fossilized arbuscular mycorrhizae are found from 460 Ma (Redecker et al, 2000), and fungi may have been present much earlier at 1430 Ma (Butterfield, 2005). The presence of Precambrian Lichens is evidentially ambiguous but nevertheless widely postulated (Selosse & Tacon, 1998, Retallack, 1994, Lenton & Watson 2004). The idea is given weight by strong evidence for terrestrial photosynthesizing microbial biota from about 1400 Ma (Horodyski & Knauth, 1994), as well as molecular clock inferences for the presence of lichen-forming fungi from about 1000 Ma (Heckman et al, 2001).

Lichens exhibit a vastly greater ecological range than any other form of macroscopic photosynthetic life. There is no biome on Earth (i.e. including the Arctic) that lacks colonization by lichens. In respect of virtually all physiologically limiting environmental variables the eco-physiological range of lichens significantly exceeds that of vascular plants or mosses (e.g. Nash, 1996). Antarctic lichens have a temperature range in the region of −2 − 14°C (Gannutz, 1967) and are capable of surviving prolonged periods below −13°C (Clark et al, 2001). Heat-adapted lichen species have a similar robustness in the opposite direction, with desert lichens capable of surviving up to 55°C (Kershaw, (1985). Thus, the extremes of temperature, water potential and availability of nutrients and rock substrate that characterize the Cryogenian do not preclude the colonization of the land surface by lichens during intervals of habitability, indeed if anything they increase the strength of the case for their presence (in comparison to that for other forms of photosynthetic life). Nor is it implausible that lichen spores survived non-habitable intervals during the glacial period, for instance lichen spores have been shown to survive in space (Sancho et al, 2007). By contrast, the optimal temperature for photosynthesis in a typical vascular plant is typically in the region of 15 − 20°C, with a far sharper cut-off outside this optimum than lichens (Hatfield & Prueger, 2015).

### D. Key premises

Life can be conceived of as introducing a unique variability to the Earth system (Vernadsky, 1926), on the basis of which we take the qualitative difference in SBE susceptibility between the Earth system before and after the Precambrian-Cambrian boundary, as plausibly biological in origin. The above discussion serves to justify the following basic premises:

1. If any macroscopic photosynthetic land life was present around and immediately after the Cryogenian period, it was likely to be lichens.
2. Any Precambrian lichen-based terrestrial biosphere likely exerted a comparable silicate weathering enhancement to the subsequent Phanerozoic vascular plant-based terrestrial biosphere, possibly a greater one.
3. Any such weathering enhancement factor was likely sustained over a greater physiological range when resulting from lichen activity that other photosynthetic life.

This leads us to the key premise motivating this work: The Precambrian terrestrial biosphere enhanced silicate weathering induced *CO*_2_ draw-down over a significantly wider temperature range than the Phanerozoic terrestrial biosphere, but the overall magnitude of this enhancement was comparable. On the basis of this supposition, we now proceed to model the likely impact of biotic enhancement of silicate weathering on the susceptibility to a SBE glaciation.

## 2. A conceptual model

The global silicate weathering flux *W* can be written:

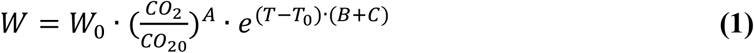

Where *W*_0_ = 6.65 × 10^12^*molCyr*^−1^ denotes a baseline silicate weathering flux value, *A* = 0.25, *B* = 0.056 and *C* = 0.017 respectively denote sensitivity to *CO*_2_ (measured in parts per million and normalized to pre-industrial levels *CO*_20_ = 280*ppm*), Arrhenius temperature kinetic sensitivity, and sensitivity to changes in runoff and hydrology coupled to temperature (Walker et al, 1981). The relationship between silicate weathering and the long-term dynamics of *CO*_2_ can be expressed in its simplest possible form as:

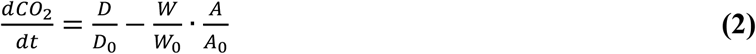

Where 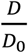 and 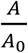 respectively denote the normalized values of the tectonic outgassing rate and the land surface area available for silicate weathering (relative to present day in each case), and *CO*_2_ can therefore be computed as simply 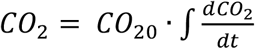 to convert to units of parts per million (ppm). Schwartzman (Schwartzman & Volk, 1989, Schwartzman 1999, 2017) devised a formulation by which biotic enhancement of weathering could be estimated without explicit parameterization of the hypothetical weathering rate on an abiotic Earth. Setting (2) to steady state, rearranging and labelling 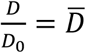 and 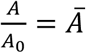, one can write the relative biotic enhancement of the global silicate weathering flux *BEW* as:

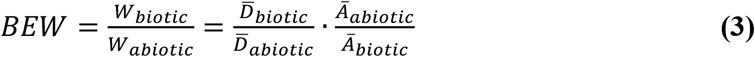

Schwartzman’s formulation entails substituting (1) into (3) then solving for 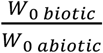, i.e. the present day biotic enhancement of weathering:

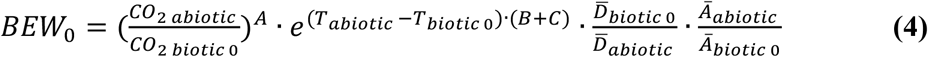

Where for each term, the zero subscript refers to the boundary conditions necessary for steady state in (2) at present day. The *CO*_2_ and temperature levels given refer to those necessary for steady state in (2) on a planet Earth with and without life. An advantage (Schwartzman, 2017) of (4) is that one can then express the biotic enhancement of weathering relative to its present-day magnitude 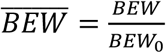, which allows one to cancel out the abiotic terms. This cancellation entails the assumption that an abiotic planet Earth would have been significantly less variable over its history than the real Earth, such that for each of the above variables; i.e. for *X* = (*T*, *W*, *D*, *A*, *CO*_2_), *X*_*abiotic*_ ≈ *X*_*abiotic* 0_. Thus, Schwartzman’s measure of biological enhancement of silicate weathering relative to that at present day is:

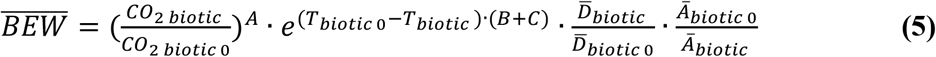

The “runaway” nature of the positive feedback between increasing ice cover, increasing planetary albedo and decreasing planetary temperature, means that equilibration of the hydrological cycle processes relevant to a snowball Earth occur over timescales of the order of ~20 years (Pierrehumbert, 2005); i.e. far shorter than the thousand-year timescales relevant to the silicate weathering “thermostat” (Walker et al, 1981, Berner et al, 1983). Consequently, in a simplistic model of geologic timescale changes, it is acceptable to discretize the changes in ice-cover and planetary albedo associated with glacial entry and exit (North, 1975). We therefore set planetary albedo *α* constant outside the temperature region *T*_*out*_ ≤ *T* ≤ *T*_*in*_ corresponding to the ice albedo instability, whilst if temperature falls below *T*_*in*_ from a non-glaciated state the albedo instantaneously jumps to a maximal “snowball Earth” value *α*_*sbe*_ = 0.7, whereas if temperature enters this unstable region from within a snowball, albedo jumps in the opposite direction to the lower value *α*_*non sbe*_ = 0.3 (Caldeira & Kasting, 1992, Pierrehumbert et al, 2005). The “time derivative” of planetary albedo therefore comprises one of three distinct values, depending on temperature’s current value and its trajectory:

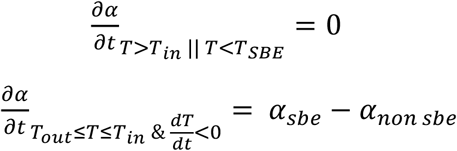

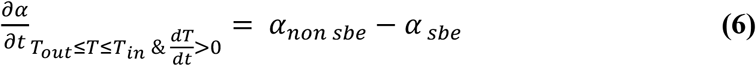

Thus, once temperature falls below a critical threshold *T*_*IN*_, a positive feedback between increasing ice cover and decreasing planetary temperature causes planetary albedo to (geologically) jump from a low value *α*_*non sbe*_ characteristic of the non-snowball Earth state, to a higher value *α*_*sbe*_ characteristic of the equilibrium state at the end of this ice albedo feedback, whilst temperature correspondingly jumps to a minimum value *T*_*SBE*_ characteristic of global scale glaciation. The analogous process occurs in the opposite direction once temperature rises above a deglaciation threshold *T*_*out*_.

The equilibrium radiative temperature of a “blackbody” planet Earth is given (in Kelvin) by:

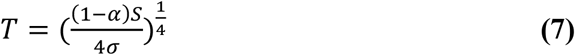

Where *σ* = 5.67367 × 10^−8^*Wm*^−2^ is the Stefan Boltzmann constant (relating the energy radiated by a surface to the fourth power of its absolute temperature), the factor 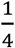 is the ratio between the area of the disc intercepting incoming solar radiation and the sphere emitting it, *S* is incoming solar radiation. For the simple first-order arguments we wish to make here, we use a linearized approximation to the radiative component of planetary equilibrium temperature, which gives the actual temperature (in Celsius) when added to a simple formulation of the climate sensitivity (North et al, 1981, North, 1990):

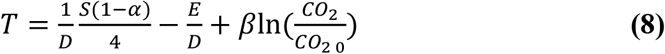

Where *E* = 203.3*Wm*^−2^*C*^−1^, and *D* = 2.09*Wm*^−2^*C*^−1^ are derived from the results of complex climate models and represent the effect of water vapour and infrared absorbing gases, and *β* = 4.33 corresponds to an equilibrium climate sensitivity of *β* ln(2) = 3.002°C per doubling of *CO*_2_, which is compatible with sensitivity analyses performed on ensembles of high resolution climate models (Cox et al, 2018). The solar luminosity flux can be related to the time of interest by (Gough, 1981, Kasting, 2005):

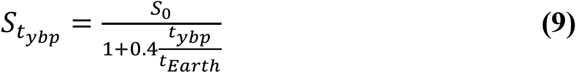

Where *S*_0_ = 1368*Wm*^−2^ is the current solar luminosity flux, *t*_*ybp*_ is the time in billions of years before present day and *t*_*Earth*_ = 4.6 is the age of the Earth in billions of years. (For the simulations shown below we set *t*_*ybp*_ = 0.7). Our objective here is simply to derive a simple relationship between the temperature thresholds relevant to the unstable region of the ice-albedo feedback *T*_*out*_ ≤ *T* ≤ *T*_*in*_ and *CO*_2_, which is satisfactory to inform upon our arguments about biological weathering enhancement. Substituting, we have the *CO*_2_- level corresponding to a prescribed glaciation entry threshold *T*_*in*_, at a given point in Earth history:

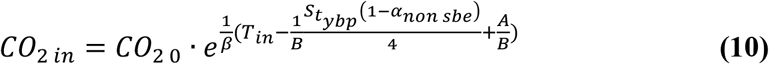

In the first instance, we can examine plausible levels of biological weathering enhancement relevant to the snowball Earth problem by directly substituting *CO*_2 *in*_ and *T*_*in*_ into (5), and taking the biotic terms as their present-day baseline values:

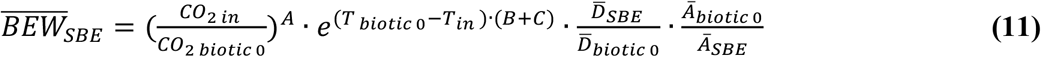

Then, setting (2) to steady state and substituting in (1), the temperature *T*_*in*_ and *CO*_2 *in*_ combination necessary for the system to move into the unstable region of the ice-albedo feedback and trigger a global scale glaciation event must conform to:

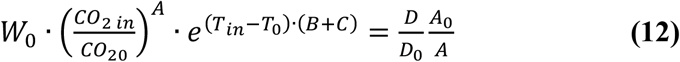

We can also write the magnitude *ɛ* of the enhancement of the silicate weathering flux as a function of a maximal enhancement factor *ɛ*_*max*_, the deviation that temperature *T* exhibits from the optimum *T*_*opt*_ for the growth of the biota responsible for this weathering enhancement (for the sake of parsimony we assume both positive and negative temperature deviation from optimum are equally detrimental) and the sensitivity *k* of biological weathering enhancement to this temperature deviation:

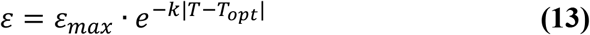

In the interests of parsimony and clarity we approximate *T*_*opt*_ ≈ 15°C as a broad average for optimal productivity and take *k* as the parameter encompassing the nuances of the distinct ecophysiology of different species. As discussed above, our central hypothesis is that the Neoproterozoic Earth system, in which any biotic weathering rate enhancement was likely driven by lichens and bacteria on the land surface, was characterized by a much lower temperature sensitivity than the Earth system of the Phanerozoic:

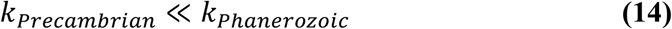

We begin by rewriting Schwartzman’s metric in (3)-(5) using a modified silicate weathering flux, i.e. multiplying (1) by (10):

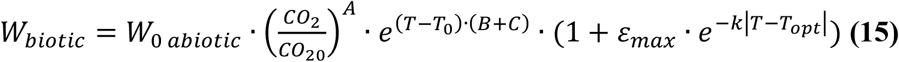

A crucial theoretical consideration concerns the relationship between biotic enhancement of silicate weathering and susceptibility to snowball Earth. If life has changed the susceptibility that Earth exhibits to snowball Earth (for a given set of abiotic boundary conditions), then the conditions describing the entry point to a snowball Earth are slightly modified to:

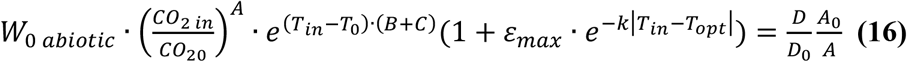

Where obviously *ɛ*_*max*_ = 0 causes (16) to revert to the same form as (12). Thus far this is merely a relabelling exercise. But suppose that we set *T*_*opt*_ ≈ *T*_0_ ≈ 15°C (i.e. assume that the optimum temperature for biotic weathering enhancement does not differ significantly from the current planetary average). Then when *T* = *T*_0_ (12) is equal to:

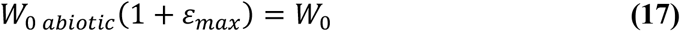

(where *W*_0_ is the above actual value of the silicate weathering flux used as a baseline in simple climate models). We can parameterize:

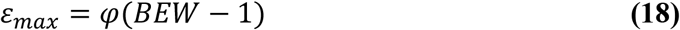

(where 2 ≤ *BEW* ≤ 5.4 denotes bulk enhancement of chemical weathering by life, estimated within various existing modelling and empirical studies, discussed in detail by Schwartzman (Schwartzman, 2017), and *φ* is a tuneable parameter (see figure 2)). Which therefore allows us to estimate the baseline scaling factor for the silicate weathering flux on a hypothetical abiotic planet:

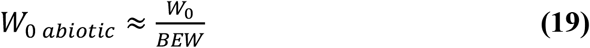

Formally therefore, the crux of the hypothesis on which we focus here is that:

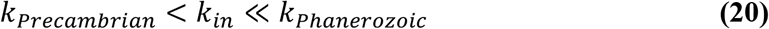

Which amounts to the idea that biological evolution drove changes in the temperature sensitivity of the terrestrial biosphere that caused, under comparable abiotic boundary conditions, Earth to cease to become susceptible to a snowball Earth event after the colonization of the land by vascular plants, and the rise of complex soil surfaces etc, into the Phanerozoic. Using (12) and (16) the following state of affairs describes necessary and sufficient conditions for a difference between the two formulations of biotic enhancement of silicate weathering in respect of susceptibility to SBE inception:

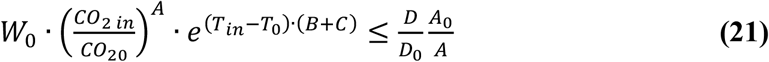

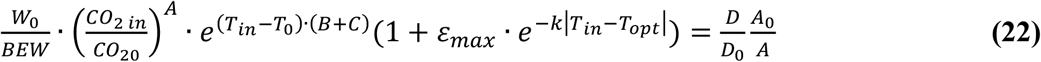

If (21) is an equality (i.e. the system nearly enters the unstable region of the ice-albedo feedback without biotic weathering enhancement, but fails to do so), (21) and (22) can be equated and we can divide through by the left hand side of (21) and solve for the maximum temperature sensitivity *k*_*max*_ at which this “amplification” effect might make a difference:

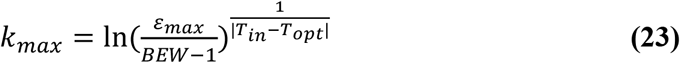

Which obviously requires *ɛ*_*max*_ > *BEW* − 1 for non-zero *k*. In summary, we contend that biological evolution has introduced a variability into *k*_*in*_ during Earth’s history, which has resulted in the weathering system coming into equilibrium with different *CO*_2 *in*_, *T*_*in*_ combinations at distinct points in time.

## 3. Results

Figure 1 depicts example simulations, figures 2 and 3 sensitivity analysis to key parameters. In light of the above discussion regarding the poor constraints on parameters within even the most sophisticated high-resolution models, we emphasize that the results given here should be viewed as ultimately qualitative and intended to emphasize a key un-investigated quantity. Figure 1 shows the difference between weathering and the product of degassing rate and weatherable continental area, at steady state, at a proscribed value (left hand columns) for the temperature *T*_*in*_ at which a runaway ice-albedo feedback to global scale glaciation occurs.

**Figure 1:**
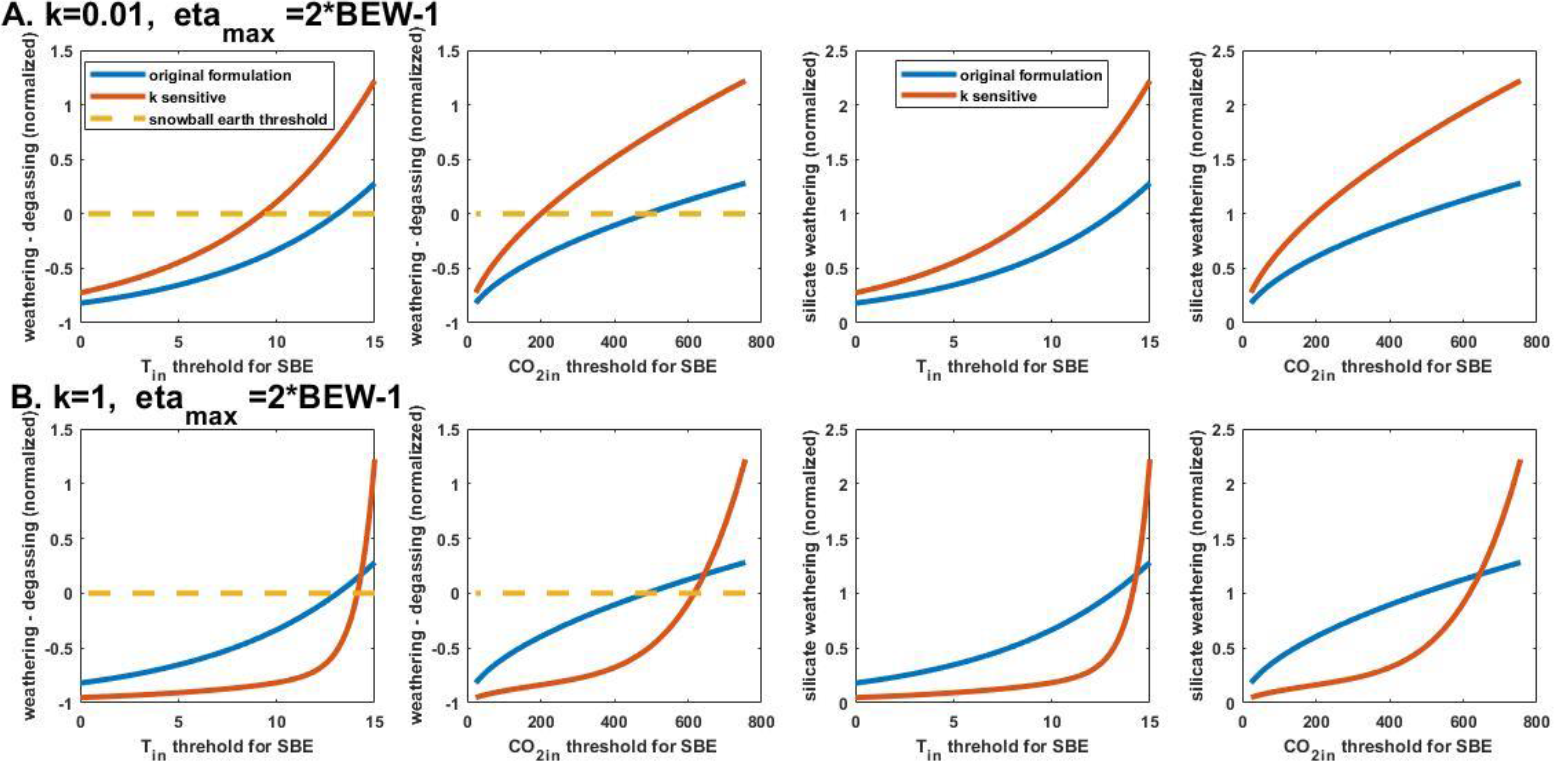
Example steady state results given different formulations for biotic enhancement of silicate weathering. The proximity of different steady state results to SBE initiation assuming biotic weathering enhancement of the form given by equation s(12) (blue lines) and by equation (16) (red lines). In each row, the left-most column shows the steady mass balances 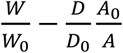 for the formulations of weathering *W* given in equation (12) (blue lines) and (16) (red lines), plotted against *T*_*in*_ (leftmost column), and against *CO*_2 *in*_ (second from left). The next two columns in each row show the corresponding magnitudes of the silicate weathering fluxes, again plotted against *CO*_2 *in*_ (rightmost column) and *T*_*in*_ (second from right). Row A (above) shows a temperature-sensitive biotic enhancement factor with a relatively low sensitivity to deviation from optimal temperature *k* = 0.01, *φ* = 2, row B (below) shows a relatively high sensitivity *k* = 1, *φ* = 2. The dashed line shows the threshold that must be crossed for biotic weathering to further decrease the temperature from the steady state depicted, thus implicitly trigger a snowball Earth.

**Figure 2:**
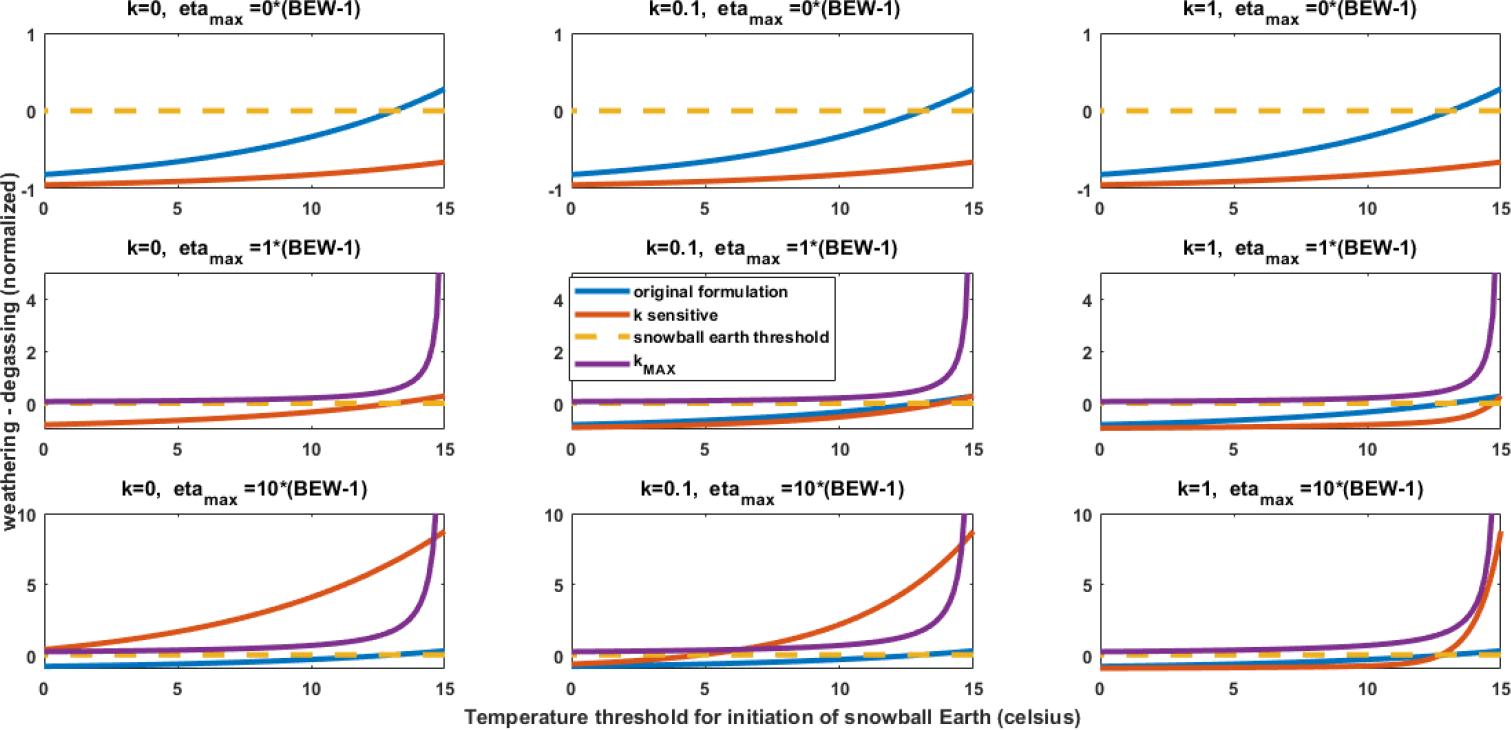
Sensitivity analysis focusing on parameters potentially influenced by directional changes in the physiology of the terrestrial biosphere. Different rows show different values for the proportionality constant *φ* (see equation (18)) relating the parameter *ɛ*_*max*_ to (the empirically constrained) bulk enhancement of chemical weathering *BEW*. As shown in equation (18), *ɛ*_*max*_ dictates the magnitude of a theoretical maximum biologically enhanced weathering rate, as opposed to *k* which dictates the sensitivity of biotic enhancement of silicate weathering to temperature deviating from the biological optimum *T*_*opt*_ and different values of which are shown across the three columns. Individual graphs and axis labels are identical to top row of figure 1. Also depicted is the hypothetical maximum value of the biotic-weathering temperature sensitivity *k*_*max*_ (equation (23)). If *ɛ*_*max*_ is zero (top row) then active biotically enhanced weathering as we have formulated it here is impossible and reverts to the abiotic rate. Once active weathering enhancement by land life occurs (middle and lower rows), biotic initiation of glaciation is possible, but (as described in the main text), as the biota responsible for this weathering enhancement become increasingly temperature sensitive (lower row, left to right), such initiation becomes less probable. Note the difference in scale on the Y-axis of different rows.

**Figure 3:**
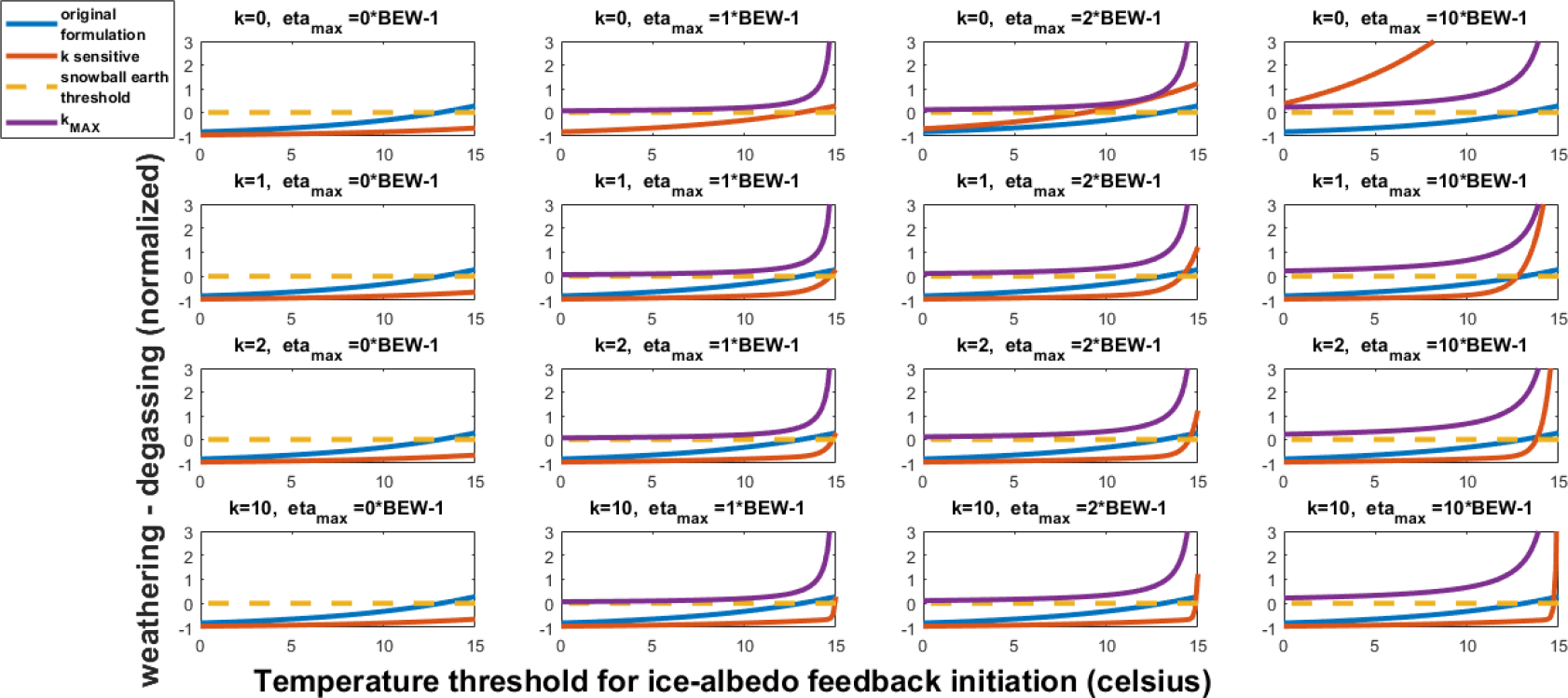
Expanded sensitivity analysis. As figure 2a, but analysis extended over a wider range of parameter values, illustrating the generality of the conclusions.

This proscribed value for *T*_*in*_ determines (by equation (10)) a corresponding greenhouse threshold for glacial entry *CO*_2 *in*_ (X-variable in the right-hand columns). The red line corresponds to temperature-sensitive biotic silicate weathering enhancement corresponding to equation (16), the blue to Schwartzman’s (1999, 2017) original, temperature-“insensitive” (beyond direction kinetic dependence in the baseline reaction rate) corresponding to equation (12). Note that the system becomes less meaningful the further the steady state is from the dashed line, because this corresponds to increasing deviation from steady state. The key quantity of interest is the value of *T*_*in*_ at which this threshold (i.e. a triggering of a SBE event) is crossed. A comparison between the red lines in the left column of the upper and lower panels illustrates that at a lower value of *k* (i.e. lower sensitivity of biotic weathering enhancement to deviation from optimal temperature) reduces the temperature threshold at which a SBE event is induced under equivalent boundary conditions.

This basic idea is robust to changes in the value of *k* used, as well as to changes in *φ* (see equation (18)), which measures the sensitivity between the *ɛ*_*max*_ parameter in equation (16) and the baseline biotic enhancement of weathering *BEW* factor taken from experiments (Schartzman, 2017). A large value of *φ* corresponds to a larger bulk value for the biotically enhanced weathering flux (across all temperatures, regardless of the biota’s sensitivity to deviation from optimum), thus effectively flattens out the curve of the relationship between *T*_*in*_ and the SBE entry threshold for a given value of sensitivity *k*. (This is illustrated by figures 2 and 3, which show results of the form of the left hand column of figure 1, across a wider range of values for these parameters). Also depicted in figures 2 and 3 (purple line) is the threshold value for the temperature sensitivity *k*_*max*_ (equation (23)), which imposes a constraint on how high this sensitivity can become whilst biotically enhanced silicate weathering rate still exceeds the baseline abiotic rate (and exhibits an asymptotic decline with *T*_*in*_).

## 4. Conclusions

As noted above, our intention here is to highlight a key unconstrained relationship that potentially connects biology with a central outstanding problem in climate science. Our key result is the comparison between relatively high and relatively low temperature sensitivities in the biotic weathering enhancement factor (upper and lower rows, figure 1). If biotic enhancement of silicate weathering is driven by a highly temperature sensitive biosphere, contribution of the biota to the initiation of a snowball Earth is only feasible at a relatively high value of *T*_*in*_. Conversely, if such enhancement is relatively temperature insensitive, biology may conceivably increase the likelihood of SBE initiation even when so-doing necessitates tolerance of a relatively low value of *T*_*in*_. The key proposition we make is therefore that any such biological “amplification” of SBE susceptibility was more probable in a lichen-dominated Precambrian Earth system than a Moss and (later) plant-dominated Phanerozoic Earth system.

In essence, we suggest that the upper row of figure 1 is qualitatively representative of the Precambrian biosphere and the lower row of that of the Phanerozoic biosphere. Our core idea is relevant to the snowball Earth problem if (for the parameter choices used), *T*_*in*_ ≤ 10°C, because then only the low-temperature-sensitivity “Precambrian” system can induce a SBE event. Crucially, we suggest that at certain points in Earth’s history, the baseline silicate weathering flux (just) self-limited on the approach to a snowball Earth (via the temperature dependence of the weathering reaction), but that biology evolved to overcome this by compensating for the temperature decrease (due to the intensity of the selection pressure to extract nutrients from rocks etc). This may have been sufficient to tip the system into a snowball Earth state; in a context in which it would otherwise have remained merely at the edge of this state.

Clearly it is not possible to reconstruct the species composition by mass, land surface coverage and temperature tolerances of any (hypothetical) Precambrian land biota. We justify our qualitative statement about temperature tolerance purely on the basis of the unique eco-physiological robustness of lichens. Whilst not wishing to ignore the existence of cold and heat-adapted vascular plants, it is clear that evolutionary adaptation is capable of pushing lichen physiology to a far wider temperature range than vascular plant physiology. Thus, a terrestrial biosphere dominated by vascular plants will necessarily have a far narrower window of tolerance for deviation from the optimal temperature for growth and photosynthetic carbon fixation. This statement will still hold regardless of the uncertainties in estimating such physiological properties in ancient land life.

Previous work has linked lichen-induced weathering to snowball Earth (Lenton & Watson, 2004, Mills et al, 2011, 2017). But these studies have focused on the connection between 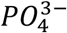 extraction from the land surface in relation to connections between the marine phosphate reservoir and ancient oxygen levels (Lenton & Watson, 2004), or on changes in degassing rate and their relationship to glacial inception (Mills et al, 2017). The temperature sensitivity of the silicate weathering flux (Walker et al, 1981) implies a self-limitation process for any (biotically or abiotically) weathering-induced glaciation: As *T*_*in*_ is approached, weathering necessarily declines, slowing the rate of any such approach. When we use the phrase “biological amplifier” above, we mean to suggest that biotic enhancement of weathering may potentially have compensated for such a decline and permitted entry to the unstable region of the ice-albedo feedback. The boundary conditions relative to which this is possible (temperature, degassing rate, land surface area, hydrological cycle feedbacks) are obviously crucial additional parameters that must be constrained in order to assess whether this is plausible (and are obviously not something we have addressed here). Experiments cross referencing the existing estimates (Schwartzman, 2017) for various forms of biological enhancement of weathering against temperature extremes are urgently needed, so as to determine the sensitivity to temperature of biotic weathering enhancement. If any such sensitivity is significant and differs between lichens and other photoautotrophs and significant weathering-enhancing organisms, then we propose that our arguments here highlight a potentially important missing parameter that should be included in high-resolution models of snowball Earth.

